# Improved DNA based storage capacity and fidelity using composite DNA letters

**DOI:** 10.1101/433524

**Authors:** Leon Anavy, Inbal Vaknin, Orna Atar, Roee Amit, Zohar Yakhini

## Abstract

DNA, with its remarkable density and long-term stability, is an appealing potential next generation data storage medium, most notably for long-term archiving. Megabyte scale DNA based storage was first reported in 2012. The Shannon information capacity of DNA was recently demonstrated, using fountain codes, to be ∼1.57 bit per synthesized position. However, synthesis and sequencing technologies process multiple nominally identical molecules in parallel, leading to significant information redundancies. We introduce composite DNA alphabets, using mixed DNA base types, to leverage this redundancy, enabling higher density. We develop encoding and decoding for composite DNA based storage, including error correction. Using current DNA synthesis technologies, we code 6.4 Megabyte data into composite DNA, achieving ∼25% increase in capacity as compared to literature. We further demonstrate, on smaller scales, how flexible synthesis leads to 2.7 fold increased capacity per synthesized position. Composite DNA can thus reduce costs for DNA based storage and can also serve in other applications.

---

DNA based data storage systems are particularly appealing due to the high information capacity of DNA compared to current state of the art storage media. Storing digital information on DNA involves encoding the information into a sequence over the DNA alphabet (i.e. “A”, “C”, “G” and “T”), producing synthetic DNA molecules with the desired sequence, and storing the synthetic biological material. Reading the stored information requires sequencing of the DNA molecules and decoding the resulting sequence to obtain the original digital information. Here we demonstrate a complete DNA based storage system with higher information density then previously reported. We introduce an approach that leverages inherent properties of DNA synthesis and sequencing, specifically the fact that DNA is synthesized and sequenced with high multiplicity^1,2^.

Demonstrations of DNA based storage systems^3–10^ raise unique technological and design challenges. Biochemical and technical constraints require the use of custom coding schemes to accommodate possible dropouts and common DNA synthesis and sequencing errors^6,9,11^. Random access at reduced sequencing overhead requires efficient design of large pools of mutually compatible PCR primers^7,8,10,12^. Recently, novel synthesis approaches were introduced^13,14^ that may lead to more cost-effective DNA based data storage. Other molecular biology techniques can also be utilized to implement DNA based storage^15^ Modern DNA synthesis technology, which is based on phosphoramidite chemistry^1^, yields high numbers of molecules for each of the designed DNA sequences^16^. Oligonucleotide multiplicity, which is an important inherent property of current DNA synthesis and sequencing technologies, is not exploited by the aforementioned work. In this article we introduce composite DNA letters, constructs that utilize this multiplicity, and thereby increase the information capacity per synthesized position above the strict, single molecule, theoretical limit of 2 bits per synthesized position.

A composite DNA letter is a representation of a position in a sequence that constitutes a mixture of all four standard DNA nucleotides in a specified pre-determined ratio. As we describe herein, composite DNA letters form the basis to a DNA synthesis approach that trades sequence multiplicity for increased complexity of the synthesized DNA. This increased complexity effectively extends the available alphabet and therefore allows higher data capacity per synthesized position. Mixtures of standard nucleotides and degenerate bases were proposed as enhancing components in several biotechnology contexts. In the early days of DNA sequencing by hybridization, degenerate and semi-degenerate bases were proposed as wildcards for increasing the fidelity of the system^17–19^. Next generation DNA sequencing by single nucleotide addition have higher quality and capacity when using degenerate base addition together with error correction approaches, as was recently demonstrated^20^. Degenerate synthetic DNA sequences were also used to explore large sequence spaces and map the phenotypic landscape of complex molecular mechanisms^21^. Similar increased complexity can also be achieved, to a limited extent, by introducing synthetic orthogonal nucleotide pairs^22^.

We demonstrate an implementation of a complete large-scale composite DNA storage system using commercially available DNA synthesis and sequencing technologies and thereby demonstrate, in practice, information capacity which is superior to the previously proposed systems. Our improved capacity system implements an error correction scheme that combines an adaptation of the previously reported fountain code^9^ and an efficient Reed-Solomon coding over the appropriate finite fields, depending on the composite system used. We used this composite DNA coding system to repeat the original DNA fountain experiment^9^ and demonstrate a 24% increased capacity per synthesized position. We also stored a compressed file containing an html version of the Bible in both Hebrew and English taken from the Mamre Institute^23^. We analyze performance as a function of process characteristics to better evaluate challenges and potential improvements of composite DNA based data storage. On a small scale we demonstrate a molecular implementation at an even greater capacity of 4.3 bits per position.

## Results

### Composite DNA letters extend the available alphabet and increase data capacity

A composite DNA letter is a representation of a position in a sequence that constitutes a mixture of all four standard DNA nucleotides in a specified pre-determined ratio σ = (σ_*A*_, σ_*C*_, σ_*G*_, σ_*T*_) where *k* = σ_*A*_ + σ_*C*_ + σ_*G*_ + σ_*T*_ is defined as the resolution parameter of the composite letter (See Online Methods for details). For example, σ = (1,1,2,0) represents a position in a composite DNA sequence of resolution *k* = 4 in which there is a 25%, 25%, 50% and 0% chance of seeing A, C, G and T respectively. Writing a composite DNA letter at a given position of a DNA sequence is equivalent to producing (synthesizing) multiple copies (oligonucleotides) of the sequence, so that in this given position the different DNA nucleotides are distributed across the synthesized copies according to the specification of σ. Reading a composite letter requires the sequencing of multiple independent molecules representing the same composite sequence and inferring the original ratio or composition from the observed base frequencies (Figure 1). Introducing composite letters extends the available alphabet and thus allows the coding of longer messages within a fixed synthesized molecule length. A composite DNA alphabet is a set of composite DNA letters, usually, but not necessarily, sharing a common resolution *k*. The full composite alphabet of resolution *k*, denoted Φ_*k*_, is the set of all σ = (σ_*A*_, σ_*C*_, σ_*G*_, σ_*T*_) so that ∑_*i*∈{*A,C,G,T*}_ σ_*i*_ = *k*. Note that 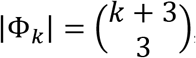, thus the composite alphabet size grows with the resolution parameter and so does the potential data capacity (Figure S1).

**Figure 1:**
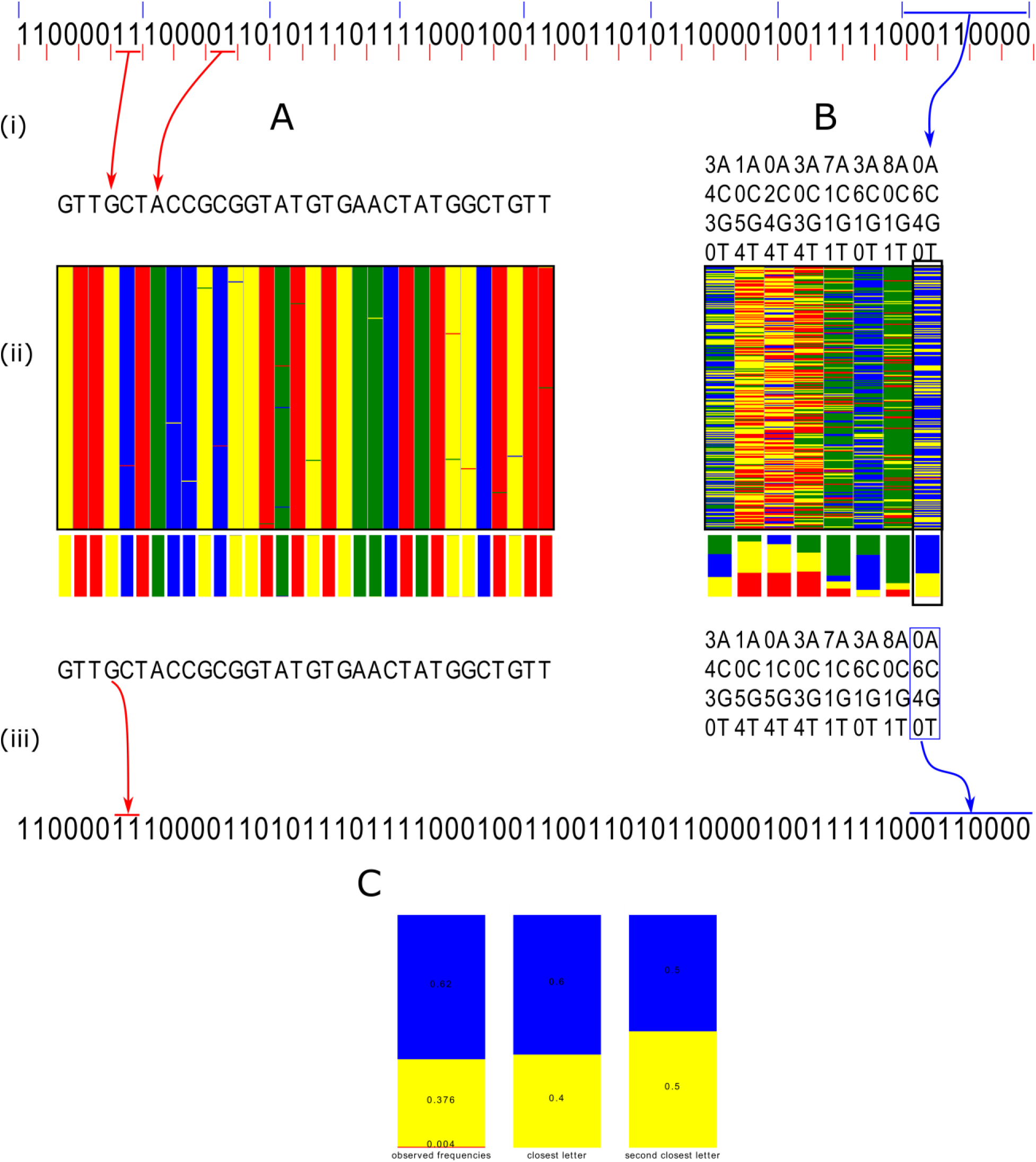
Encoding a binary message using standard and composite DNA. A binary message, depicted on top, is encoded into DNA. A. Standard DNA based storage scheme^9^. The binary message is being encoded to DNA by mapping every 2 bits (depicted by the short red separating lines) to a DNA base or synthesized position (i), the designed DNA sequence will then be synthesized and sequenced by a noisy procedure that introduces some errors (ii). The sequencing output is then used to infer the DNA composition at every position (iii). Decoding of the original message is done assuming the use of an error correcting code over the binary message. B. The same message is encoded using a composite DNA alphabet of resolution *k* = 10 by mapping every 8 bits (depicted by the blue separating lines) of the binary message to a single composite DNA position. Using sufficiently deep sequencing allows to correctly identify the original composite letters (the right most position, in a black frame, is exemplified in C) and to decode the message. The decoding also uses an error correction mechanism (Reed-Solomon over the appropriate finite field, in our implementation), over the composite alphabet. C. An example of the inference step at a single synthesized position. The observed frequencies are used to infer the source, σ = (0,6,4,0), as the closest composite letter, using KL divergence (see text and Online Methods). Note that the inference at any fixed position is affected by the sequencing depth obtained there as well as by sequencing and synthesis errors.

To read a message coded using composite DNA letters correctly we must infer the original composite letter in sufficiently many positions of the sequence from the observed reads. The sequencing readout (i.e. The observed sequencing reads) is the product of a complex process, consisting of DNA synthesis, long term storage^4,24^, sampling and DNA sequencing. The distribution of counts, for every letter in {*A, C, G, T*}, resulting from σ at depth *N* can be described by a single model in which the readout counts are multinomial:

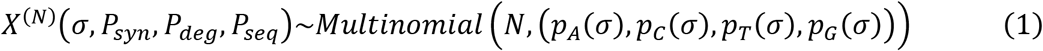

The parameters of the distribution are the designed input letter σ, the sequencing depth *N*, and the errors introduced in the synthesis, storage and sequencing steps of the process, *P_syn_, P_deg_*, and *P_seq_* (See Online Methods). While each step introduces different errors and biases, the most significant parameters that affect the readout are the sampling of molecules to be sequenced and the sequencing depth.

The sequencing readout frequencies will most likely not exactly match any letter from the original alphabet. Inference of the original letter is performed by converting the readout to a vector of base frequencies and comparing it to the base frequencies of the candidate letters in the composite alphabet. The comparison can be done, for example, using the Kullback–Leibler divergence (KL) or the *L*^1^ norm. To assess the performance of the inference step we developed a simulation model and analyzed the inference rate of the two methods on various composite alphabets (Figure S2). The KL divergence, which corresponds to a maximum likelihood estimator (See Supplementary Notes), was found, also in practice, to perform much better and was thus used in the remainder of this study, including for the molecular implementation parts (See Online Methods, Figures S3 and S10).

### Large scale implementation of a composite DNA based data storage system

To show the feasibility of the composite DNA letters concept and to demonstrate its potential for improving DNA based data archiving systems we performed a large-scale molecular implementation of a six-letter composite alphabet storage system. The system consists of using our composite letter coding approach with an adapted version of the DNA fountain system^25^ (See Online Methods) to produce a composite DNA encoding pipeline (Figure 2). We first used our system to store and successfully retrieve the same 2.12MB data file from Erlich *et al*.^25^. Our encoded DNA pool consisted of 58,000 six-letter composite oligos of length 152 nucleotides, compared to 72,000 oligos of the same length required using standard DNA, demonstrating a ∼24% increase in information capacity per synthesized position (Table 1). The six letter composite alphabet used here was Σ_6_ = {*A, C, G, T, M, K*}, where *M* = (1,1,0,0) and *K* = (0,0,1,1). Note that Σ_6_ ⊂ Φ_2_ (|Σ_6_| = 6, |Φ_2_| = 10). We also improved the error-correcting scheme of the DNA fountain code shifting the Reed-Solomon error correction from the bytes level to the DNA level (See Figure 2 and Online Methods). We further demonstrate the increased information capacity of composite DNA by encoding a bi-lingual interactive version of the bible, compressed to a 6.42MB file, using three different composite alphabets. The above six-letter alphabet Σ_6_ required 174,000 oligos, while a five-letter alphabet Σ_5_ = {*A, C, G, T, M*} ⊂ Φ_2_ required 193,000 oligos and a standard four-letter alphabet Σ_4_ = Φ_1_ required 217,000 synthetic oligos, all of the same length of 152 nucleotides (Table 1). These composite DNA oligos were synthesized by Twist Bioscience, demonstrating the first large scale composite DNA synthesis, and were used to further investigate the characteristics of composite DNA synthesis.

**Figure 2:**
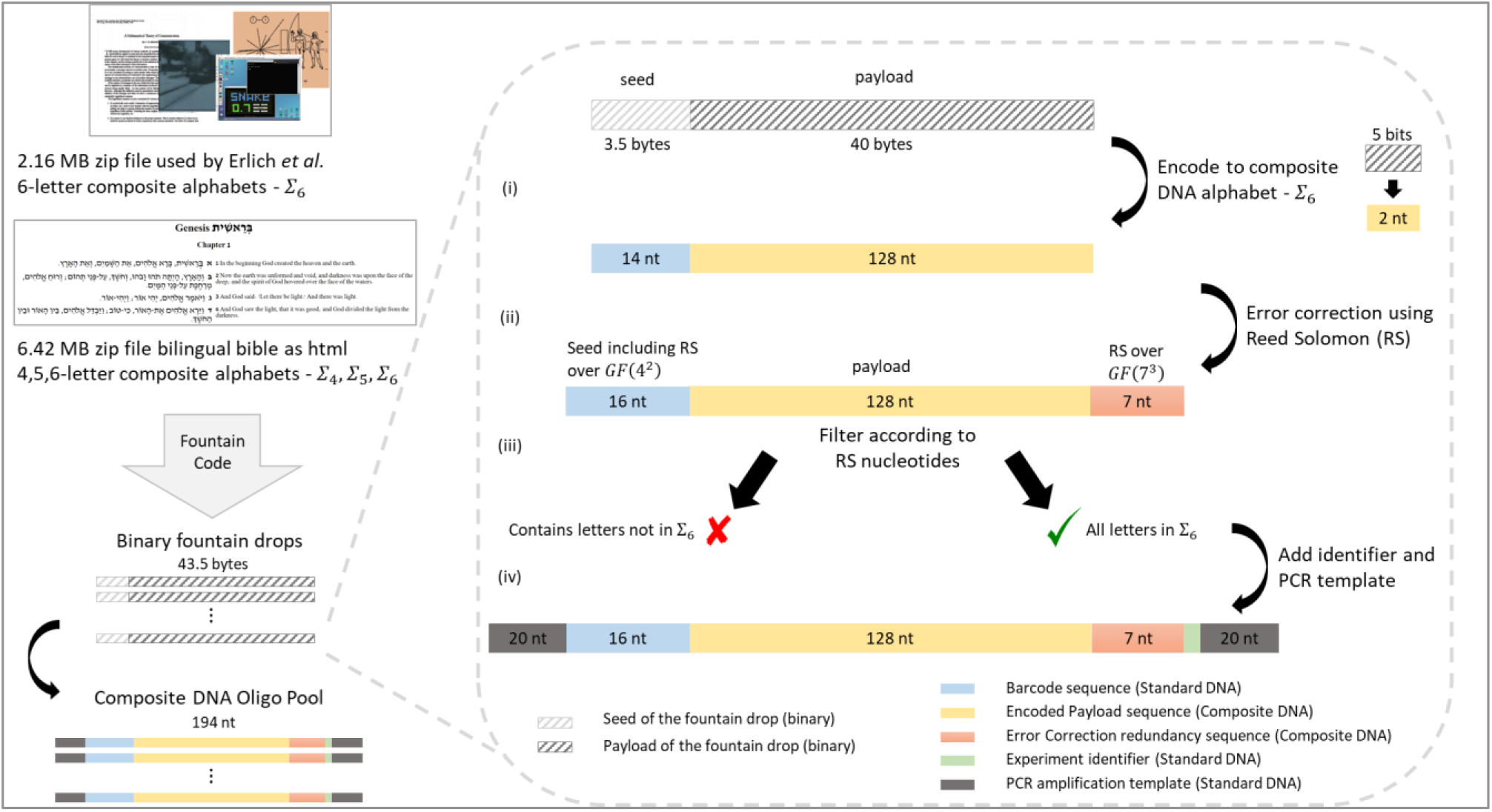
Encoding pipeline of a large scale composite DNA based data storage. A compressed input file is being processed by the fountain code to produce binary droplets. A composite DNA encoding flow is then applied on each droplet consisting of the following steps (See Online Methods for details): (i) The binary message is translated into a composite DNA sequence. The seed sequence is translated to standard DNA sequence, which will serve as a barcode for the decoding process. The payload is translated to a six-letter composite DNA alphabet (Σ_6_) in 5-bit chunks. (ii) Error correction nucleotides are added to the DNA sequence by using a systematic Reed-Solomon (RS) encoding. The barcode is encoded using RS over *GF*(2^4^) and the payload is padded and encoded using RS over *GF*(7^3^). (iii) Each encoded message is then filtered to verify that the RS redundancy letters are all from Σ_6_. (iv) Experiment identifier and amplification template sequences are appended to each valid sequence. Similar coding schemes were used for 4 and 5 letter alphabets (See Online Methods and Figures S4-S7).

**Table 1:** Comparison of DNA storage schemes using experimental results and simulations. The schemes are ordered chronologically. Information for previous studies is taken from Erlich and zielinski^25^. Information for Organick *et al*. is taken from their report. Capacity is calculated by dividing total binary input data, measured in bits, by the total number of synthesized positions (See Table S1 for details). Reed-Solomon error correction is marked as RS. Fountain code redundancy level is specified in parentheses.

| Study | Experiment | Error correction | Robustness to dropouts | Input data (Mbytes) | Total OL length (nts) | Capacity (bit/synth-position) |
| --- | --- | --- | --- | --- | --- | --- |
| Church *et al*<sup>5</sup>, 2012 |  | - | - | 0.66 | 6,313,270 | 0.83 |
| Goldman *et al*<sup>6</sup>, 2013 |  | + | Repetition (4) | 0.76 | 17,940,195 | 0.34 |
| Grass *et al*<sup>24</sup>, 2015 |  | RS | RS | 0.08 | 583,947 | 1.14 |
| Bornholt *et al*<sup>7</sup>, 2016 |  | - | repetition (1.5) | 0.05 | 18,120,000 | 0.02 |
| Erlich and zielinski<sup>25</sup>, 2017 |  | Byte level RS | Fountain (1.07) | 2.12 | 10,944,000 | 1.57 |
| Organick *et al*<sup>10</sup>, 2018 |  | Byte level RS | RS | 200.2 | ~2,000,000,000 | 1.1 |
| This work (Message from Erlich *et al*) | Composite $\Sigma_6$ | DNA level RS | Fountain (1.1) | 2.12 | 8,758,000 | 1.93 |
| This work (Bible) | Standard $\Sigma_4$ | DNA level RS | Fountain (1.08) | 6.42 | 32,767,000 | 1.57 |
| | Composite $\Sigma_5$ | DNA level RS | Fountain (1.08) | 6.42 | 29,143,000 | 1.76 |
| | Composite $\Sigma_6$ | DNA level RS | Fountain (1.08) | 6.42 | 26,274,000 | 1.96 |
| This work (Small scale) | Composite $\Phi_3$ | - | - | 22.5 bytes<br>After binary Huffman | 42 | 4.29 |

The synthesized DNA was amplified using two different primer pairs as technical repeats. We then sequenced the resulting synthetic DNA sample (100nts paired ends, Illumina HiSeq at the Technion Genome Center). Our library and reaction design allowed for separately decoding each one of the four test messages, as described above, with each of the primer pairs. In Fig 3 we describe the process of decoding the results of a sequencing reaction, performed on a synthesized composite DNA library originating from one message (in this case, the 6.4MB Bible encoded into Σ_6_ with a single pair of primers), and of inferring the underlying binary message (See Online Methods for details). Briefly, we first preprocess the raw reads by assembling paired ends reads, filtering by length and grouping by putative barcode sequence (prefixes). Next, we filter out prefixes with less than 20 associated reads generating a set of putative barcodes each associated with a group of reads. We infer the full composite oligo for each putative barcode, using KL inference. The resulting composite oligos are decoded Reed-Solomon (over *GF*(7^3^) in the case of Σ_6_) and the valid oligos are converted into binary drops. Finally, we apply a binary fountain code decoding to obtain the original message, if successful. We note (Fig 3B) that the average observed multiplicity, for each one of the inferred putative barcodes was 96 reads. Inference of the full composite oligo was only done for putative barcodes with more than 20 reads. Fig 3C and Fig 3D depict frequencies and KL inference decision boundaries (red dashed lines) for positions that are originally designed as composite. Note that individual synthesized positions are correctly inferred at an error rate of less than 10^−5^ for these filtered putative barcodes. These errors are either corrected by the Reed-Solomon code (over *GF*(7^3^) in the case of Σ_6_) or rejected by the same mechanism. Eventually, sufficiently accepted drops (172,608 for the Bible encoded into Σ_6_) made it to the last decoding step which uses an adaptation of the fountain code mechanism proposed by Erlich and Zielinski^9^ (Figure 3E). Since Φ_2_ ⊂ Φ_*k*_ for every even value of k, we can use the two composite letters from Φ_2_ (e.g K and M), to calculate correct inference rates for larger composite alphabets. In Fig 3C and 3D we further indicate, using the inner grey lines, the decision boundaries that would have been used under Φ_4_ to distinguish, e.g *K* = (0,0,2,2) from σ = (0,0,3,1). Using these decision boundaries we would have up to 7% of the positions designed as K (or M) potentially leaked to be interpreted as one of the two the neighboring composite letters. All this in the current sequencing depth. We further analyze the implication of extending the composite alphabet in Figure 4.

**Figure 3:**
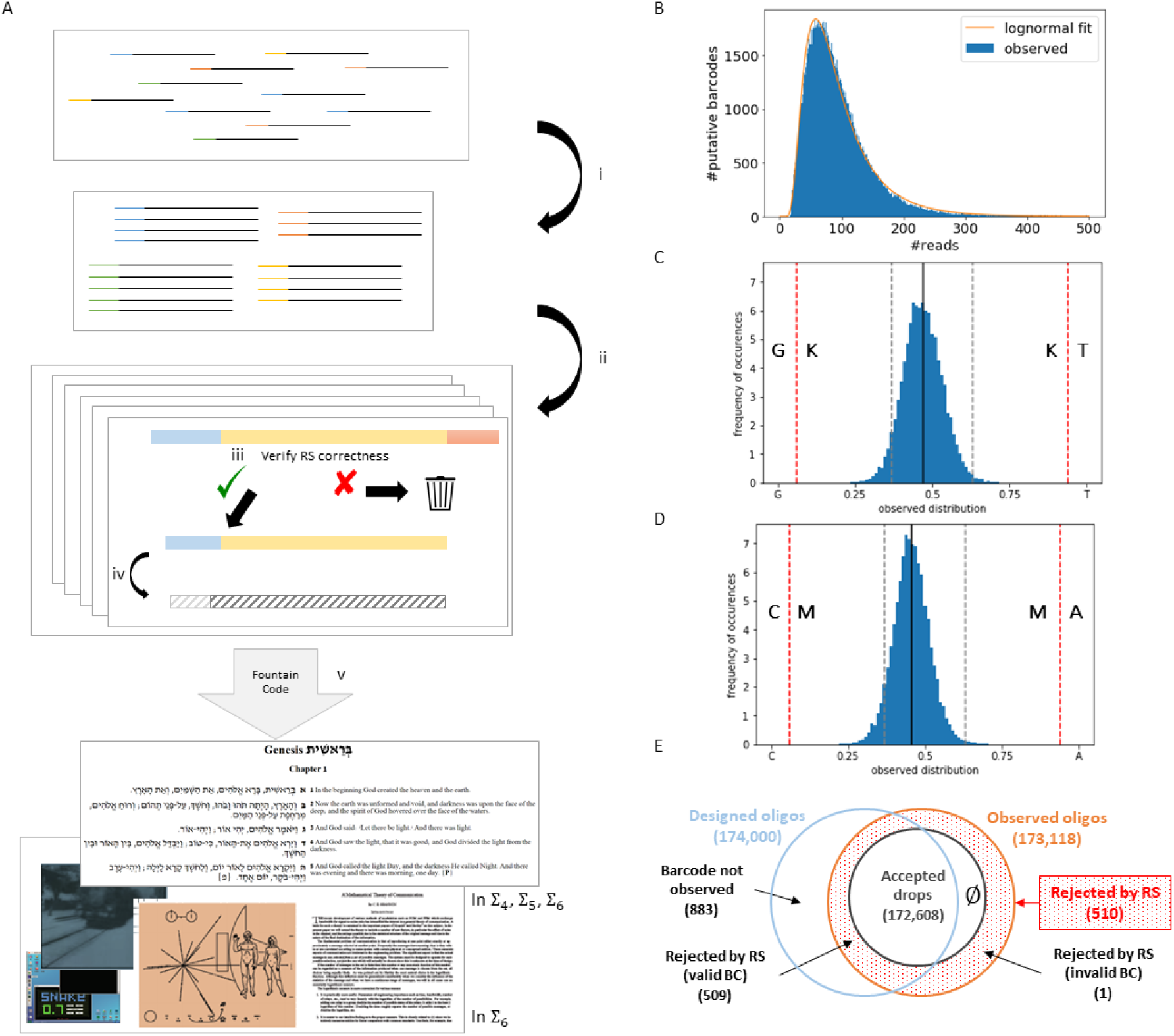
Performance of a large-scale composite DNA based storage system. Decoding a composite library to infer the original encoded message. A. The steps of the decoding process (see Online Methods): (i) Preprocess and grouping. (ii) Generation of a set of putative barcodes (iii) Inference of composite oligos using KL inference and RS error correction. (iv) Conversion into binary drops. (v) Binary fountain code decoding to obtain the original message, if successful. In panels B-E we provide some descriptive statistics related to the decoding process. Numbers indicated are for the Bible 6.4MB message encoded into Σ_6_ composite DNA. B. Number of reads associated to each 16nt prefix (putative barcode). The distribution follows a log-normal shape with a median of 81 reads and a mean of 96 reads. C-D. Distribution of base frequencies per synthesized position. For this counting we consider the positions that were designed to be composite – either K or M. KL decision boundaries also depicted, see text. E. Acceptance statistics for the designed composite oligonucleotides. D.

We observe that the mean base frequency of the composite letters K and M is slightly shifted toward G and C, respectively (Figure 4A). As an immediate result, the leakage into neighboring letters is mainly toward G and C when considering Φ_2_ decision boundaries. As expected, this leakage is diminishing as we increase sequencing depth. In particular, inference of both K and M, under hypothetical use of Φ_2_ or of Φ_4_ is perfect even at 200 reads per barcode (Figure 4B). When considering Φ_6_ we get reasonable performance at the higher depths. It is important to note that some errors in inference can be tolerated as we use a RS error correction on the complete composite oligo at the composite alphabet level.

**Figure 4:**
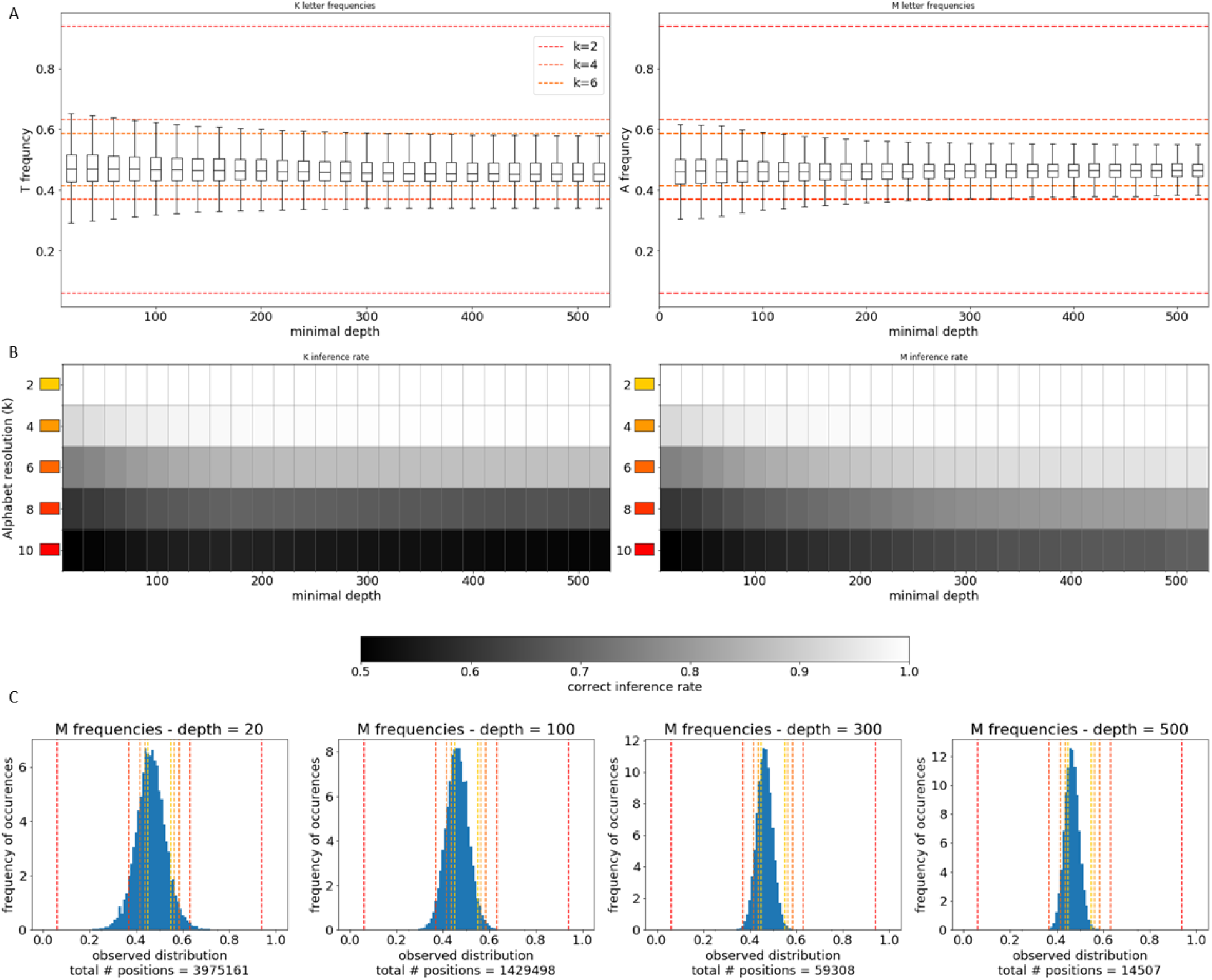
Analysis of higher resolution composite alphabets using the large-scale experiments. Sequencing depth effect on the inference of composite letters. A. Performance of the K (M) composite letter as a function of minimal sequencing depth. The boxplots depicts the distribution of observed T (A) frequencies by presenting the median, the quartiles and 1.5IQR whiskers. The dashed lines depict the decision boundaries for Φ_2_, Φ_4_ and Φ_6_. B. Successful inference rates of the composite letter K (M) over different composite alphabets as a function of sequencing depth. C. Example distribution of the T (A) frequencies for the composite letter K (M) in different sequencing depths. The dashed lines depict the decision boundaries for Φ_2_, Φ_4_, Φ_6_, Φ_8_ and Φ_10_.

### High resolution composite alphabets significantly increase data capacity

Current synthesis technology also supports the use of DNA mixtures that represent higher resolution composite alphabets, albeit on a small scale. To further explore the properties of large alphabets we encoded a short message (38 bytes in asci, 22.5 bytes after a binary Huffman compression) using composite alphabets of four different types resulting in information capacity of up to 4.29 bits per synthesized position (Table 1, Table S1 and Online Methods).

The four different alphabets used are the standard DNA alphabet, Φ_1_, the full composite alphabets Φ_2_ and Φ_3_ and an alphabet containing the 15 IUPAC letters. The input English phrase, “DNA STORAGE ROCKS!” was encoded to each of these alphabets using Huffman coding with the appropriate alphabet. The four resulting composite oligonucleotides were synthesized by IDT and sequenced by the Technion Genome Center (See Online Methods and Table S3).

First, we examined the minimal sequencing depth required to decode the message correctly for each one of the four composite alphabets. As expected, extending the alphabet by using higher resolutions requires deeper sequencing. In all four alphabets that were tested, a fully successful decoding was observed in as little as 100 reads (Figure 5A) while a near-perfect decoding was obtained with as little as 50 reads (Figure S9). In accordance with the theoretical analysis, KL inference also performed much better than *L*^1^ norm inference on the experimental data (Figure S10).

**Figure 5:**
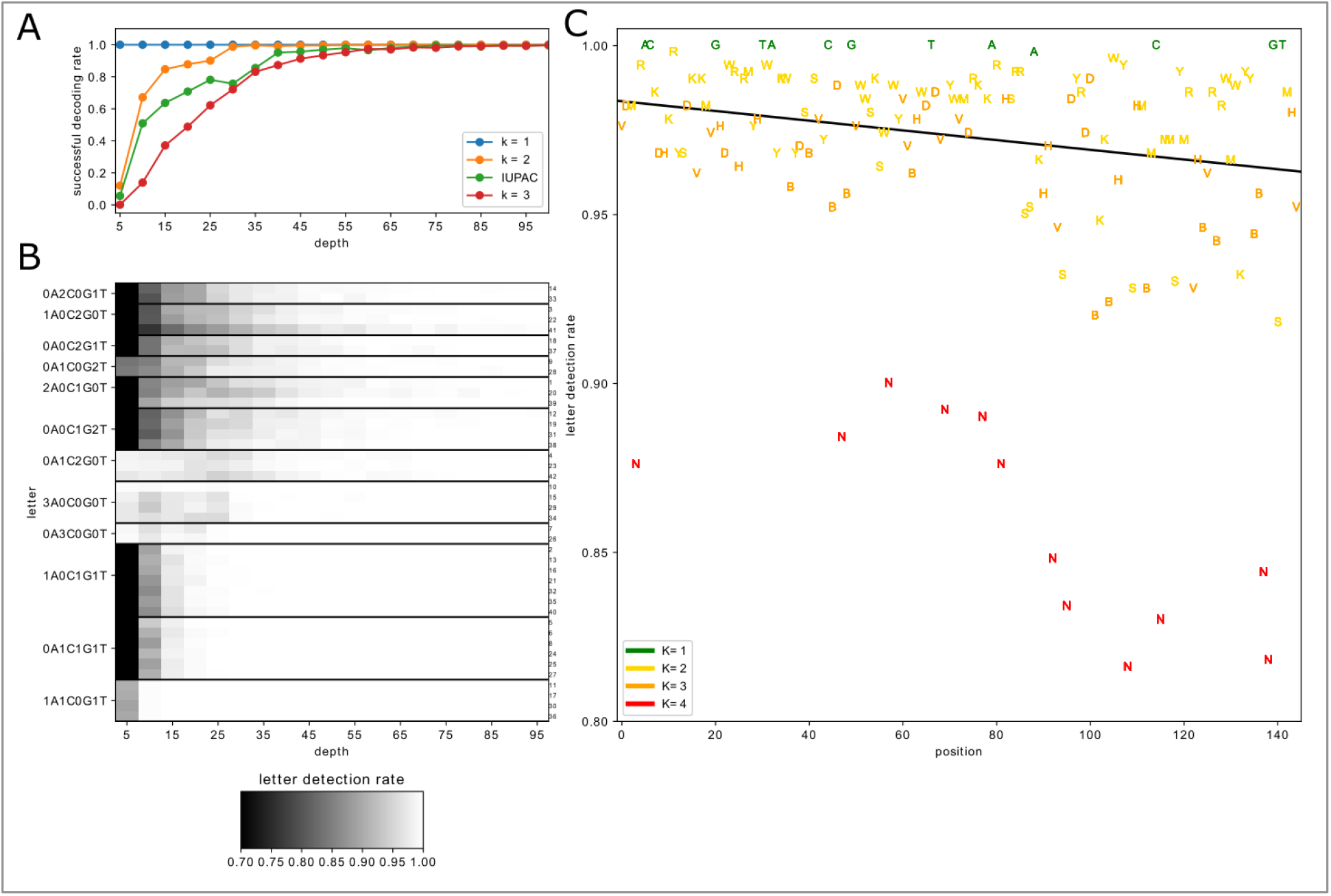
Data storage systems based on large composite alphabets. A. Successful decoding rate using KL inference as a function of sequencing depth for the four composite DNA alphabets experimentally in the molecular implementation. B. Inference rates for letters in Φ_3_ as a function of sequencing depth. The positions of the letter in the composite DNA oligo (starting from the 5’ end) are indicated on the right and the data for each letter is ordered (top to bottom) by position. C. Inference rates for the different letters of the “IUPAC” alphabet as function of position in the composite DNA sequence (starting from the 5’ end). The letters are colored according to their “native” alphabet. The black line represents a linear trend, excluding the four standard DNA letters and the single letter and “N”.

As predicted by the statistical model, some composite letters are harder to identify than others (Figure 5B). The experimental results presented above also suggest that the position of the letter in the synthesized oligo affects the identification rate. To further explore the differences between different composite letters we designed another synthetic DNA oligo containing all the equimolar letters (represented by the 15 letter IUPAC alphabet), with multiple copies of each composite letter distributed along the designed sequence (Materials and Methods, Figure S11). We examined the inference rate at a depth of 15 reads and reported the results as a function of the letter and the position in the oligo (Figure 5C). We observed a small but persistent decrease in inference rates considering the position on the synthesized oligo starting from the 5’ end.

To clarify the potential of combining large composite alphabets with high throughput synthesis using our composite DNA coding/decoding approach we calculated the capacity of large composite alphabets based storage systems (Table S1). A system using Φ_10_, which consists of 286 letters, achieves information capacity of 6.1 bits per synthesized position, which is a 3.85-fold increase over the standard DNA system.

## Discussion

In this work we presented the concept of composite DNA, which leverages properties of current DNA synthesis and sequencing processes, to potentially attain higher density DNA based storage systems. Composite DNA schemes can be combined with other approaches to increase capacity and fidelity of DNA based storage systems such as orthogonal base pair systems^22^, efficient coding^8,11,25,26^ and random access approaches^7,8,12,27,28^. Incorporating composite DNA into future DNA based storage systems will require further investment in several directions. First and foremost, any large scale implementation will require scaling up the currently limited composite DNA synthesis.

Using highly multiplexed composite DNA sequences will require better understanding of the effect of composite DNA on different chemical processes involved in DNA manipulation. Previous studies dealt with the chemical limitations of these processes either by employing strict encoding schemes^5–8,11^ or by using complex coding methodologies like DNA fountains to handle sequence dropout^25^. Employing composite DNA inherently generates balanced DNA molecules, resulting from the combinatorial space associated with every designed composite sequence. While unwanted sequences will unavoidably be part of the ensemble of synthesized molecules, the inherent independence of the different positions renders them negligible, representing an extra benefit of the composite DNA approach.

The design principles for composite DNA sequences, as well as the decoding pipeline, can be further tuned for optimal results. Mixed composite alphabets can be generated to minimize inference errors without compromising the alphabet size. Technical calibration of the actual base frequencies can be added to decoding pipeline allowing the correction of systematic synthesis biases.

Using current technologies, synthesis cost per position is ∼4 orders of magnitude larger than sequencing cost per base yielding a significant potential overall cost reduction when using composite DNA. This holds despite the increase in sequencing costs entailed by required depth.

## Online methods

### Composite DNA letters definition

A composite DNA alphabet is defined as

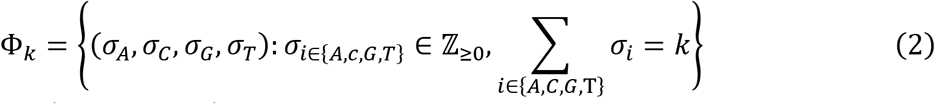

In this notation, σ = (σ_*A*_, σ_*C*_, σ_*G*_, σ_*T*_) ∈ Σ_*k*_ represents a composite letter and *k* is a tunable parameter that represents the resolution of the composite alphabet.

The size of the composite DNA alphabet grows with the resolution as follows:

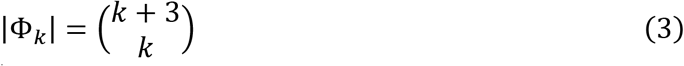

Using a naive mapping from {0,1}^∗^ to 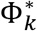 we see that the length of the encoded message decreases with the resolution of the composite alphabet: 
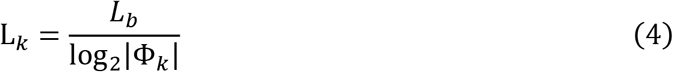
where *L*_*k*_ is the length of the message encoded using composite alphabet of resolution *k* and *L*_*b*_ is the length of the original binary message.

### A multinomial model for the composite DNA letters

The distribution of the observed count vectors is governed by the sampling process of *N* independent molecules that are sequenced and counted to generate the count vector. Given the original composite letter σ = (σ_*A*_, σ_*C*_, σ_*G*_, σ_*T*_), assuming a sequencing depth *N* (i.e. the number of independent copies sequenced), and ignoring other factors beside the sampling, the observed counts constitutes a random variable with a multinomial distribution: 
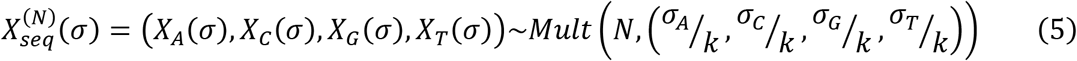
and so

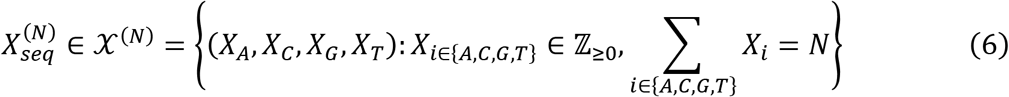

The observed read counts are, in actuality, also affected by the following parameters:

- Synthesis error rate represented by {*P*_*syn*_}_*i*→j_ = *P*(*j sy*n*thesized*|*i designed*)
- Degradation rates. {*P*_*deg*_}_*i*→j_ = *P*(*j present after storage*|*i synthesized*)
- Sequencing error rate. {*P_seq_*}_*i*→j_ = *P*(*j read*|*i present*)

Deletion and insertion events are a special class of errors in DNA synthesis and sequencing. These affect the read counts for all positions following the event position.

Assuming independence of the different error sources we can incorporate all errors into a generalized multinomial model with slightly altered probabilities:

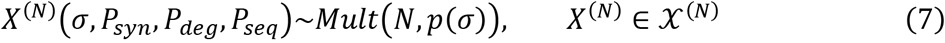

Where *p*(σ) = (*p*_*A*_(σ), *p*_*C*_(σ), *p*_*G*_(σ), *p*_*T*_(σ)) is a corrected probability vector.

### Inference of composite DNA letters

To correctly read a message coded using composite DNA letter alphabet we must infer the original composite letter σ from the observed 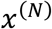. Namely, we need to define a decoding map:

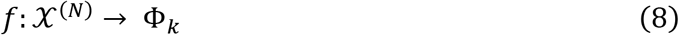

We infer the original letter from the observed vector 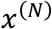 by first calculating a proportion vector 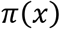:

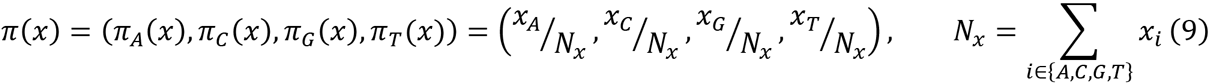

Next we use one of two mapping approaches.

*L*^*p*^ norm:

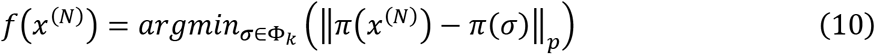

KL:

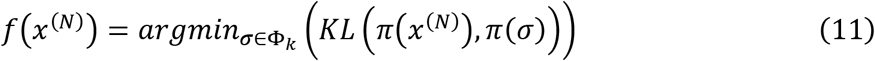

Where *KL*(*P, D*) stands for the Kullback–Leibler divergence: 
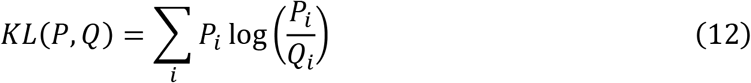
and here *i* runs over 4 letters.

When using the error-aware multinomial model, the KL approach is equivalent to a Maximum-Likelihood mapping (See supplementary text below). Since the KL measure is highly sensitive for letters on the edges of the simplex we implemented this approach using a variation of the composite alphabet in which zero entries in the probability vectors are replaced with some small value ∊.

### Simulations of composite DNA letter inference

The probability to correctly identify the original letter from the observed count vector is defined as: 
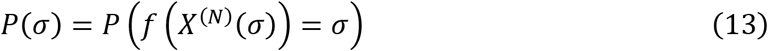
where *X*^(*N*)^ is distributed in accordance with the process parameters.

We simulated the process of DNA synthesis, storage and sequencing to examine the properties of the composite DNA inference mechanisms. For simplicity, we used a single error rate parameter. We then used the inference mechanisms described above to infer the original composite letter. For a given composite alphabet we repeated the process for *R* = 1000 times for each composite letter to estimate the inference success rate for every letter σ (Figure S2).

### Implementation of a composite DNA based storage system

We designed a composite DNA based storage system consisting of the following components:

1. DNA fountain encoding with no Reed-Solomon error correction to an extend output alphabet. We altered the previously described the DNA fountain code^9^ to support composite DNA sequences. The seed value of the DNA fountain was limited to fit in 3.5 bytes. The conversion of the binary droplet to a DNA sequence was altered so that the droplet seed, which is encoded in the first 3.5 bytes, was converted to a 14 nt standard DNA sequence acting as a barcode, and the rest of the binary sequence was converted to the desired composite DNA alphabet.
2. Addition of Reed-Solomon (RS) error correction directly to composite DNA sequence. We implemented RS codes over finite fields of various orders. Two bases of standard DNA were added to the barcode sequence by using a systematic (16,14) RS code over *GF*(2^4^). The remaining 128 nt composite sequence was padded to be 129 = 43 × 3 nt and then encoded using a (43,45) RS code over *GF*(4^3^), *GF*(5^3^) and *GF*(7^3^) for the composite alphabets Σ_4_, Σ_5_ and Σ_6_ respectively. This generated composite sequences of length 151nt. To overcome the mismatch between the six-letter alphabet Σ_6_ and the seven-letter finite field, an additional filtration step was used in which encoded oligos were included in the final set of oligos only if all the RS redundancy bases were in Σ_6_. This entailed a 13% overhead in the generation time of the final set of oligos (See Figure S4-S7).

The resulting set of composite oligos was incorporated into a constant DNA backbone containing amplification primers of 20nt on each side (we used two different sets of primers as technical repeats) and a 3nt barcode marking the experiment ID (input dataset and output alphabet). This resulted in a set of 194nt composite oligos. Combining the sets from all four experiments and two sets of primers resulted in 1.4 million composite oligos (Table S2) that were synthesized by Twist Bioscience.

The synthetic DNA library was amplified using 14 cycles of PCR and the sequenced by the Technion Genome Center. Sequencing was done on two lanes of an illumine HiSeq machine and resulted in ∼230M reads for both primer sets.

The decoding of the message was split to two steps:

1. Generation of a set of composite oligos:

a. Pre-processing of the reads including assembly of paired end reads using PEAR^29^, filtration based on length and the existence of the primer sequence and generation of eight separate sets for the different experiment.
b. Grouping of the reads according to the 16nt prefix to generate a set of putative barcodes each with an associated set of reads.
c. Filtration of the putative barcode set to contains only sequences with at least 20 reads associated to them.
d. Inference of the composite sequence by using KL inference method.
e. Decoding of the composite sequence using the appropriate RS decoder. Only error detection was performed.
2. DNA fountain decoding of the resulting set of composite oligos using the altered DNA fountain decoder.

### Experiment with large composite alphabets

We encoded a short input message (“DNA STORAGE ROCKS!”) using an encoding pipeline consisting of the following steps:

- Mapping of the message to a binary sequence using the standard ASCI code for the English language.
- Huffman coding the binary sequence into a sequence of composite DNA letters of resolution *k* using the complete Shakespeare corpus^30^ to generate the Huffman coding scheme^31^.
- To achieve equal sequence length for all designed oligos (of different resolutions k) we repeated the encoded message to fit a predetermined length of 42 bases.

This process was performed for four different resolutions *k* = 1,2,3 and a special case in which the composite alphabet consists of only equimolar combinations of bases (representing the 15 different IUPAC codes).

To calculate the information capacity of these encodings we used similar Huffman coding of the exact same message into a binary sequence. The capacity was calculated as:

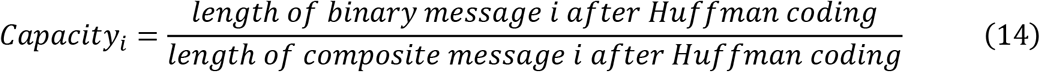

We inserted the encoded composite DNA sequence, for each of the above four configurations, into a synthetic construct containing amplification primer templates, a unique molecular identifier (UMI) and a barcode to obtain a total oligo length of 99 bases (Figure S8, Table S3).

The four designed oligonucleotides were then commercially synthesized (IDT), amplified using PCR primers from the Illumina small RNA sequencing kit and sequenced using an Illumina Mi-Seq at the Technion Genome Center.

We obtained 5,421,556 50bp paired-end reads of the four different samples. We merged the read pairs using PEAR^29^ to generate 4,855,676 reads, 95% of which with the designed length of 52 bases. We then split the reads to four different samples using the barcode value and got ∼25% of the reads in each sample.

Next, we decoded the original message using a decoding pipeline consisting of the following steps:

- Reading of the sample reads
- Filtering of the reads based on read length and removing reads containing undetermined bases (“N” output in the sequencing) and reads of length different than 52 bases.
- Inference of the composite sequence using the inference mechanisms described above.
- Decoding of the original messages using the same Huffman coding used for encoding.

For each sample we tested the ability to decode the entire message (including the repetition introduced to equalize oligo length) and also only the first occurrence of the original encoded message text.

To test for the required sequencing depth for each sample representing a specific resolution, we sampled different numbers of reads for each resolution and repeated the decoding process for each such sub-sample data. We repeated the sampling process *R* = 100 times for each sampling rate and recorded the inference rates and the overall decoding outcome for each sample.

### Error analysis for composite DNA letters

We designed a synthetic composite DNA oligo using the same overall design with the following alterations:

- The barcode and UMI were removed (unnecessary for this analysis)
- The length of the composite DNA sequence was 145 bases yielding a total oligo length of 192 bases.

The 145 composite bases consisted of all the possible pairs of composite letters. This oligo design was constructed as a de Bruijn sequence using the following methodology. A balanced circular de Bruijn sequence over an alphabet of 12 letters composed of the eleven composite letters (15 IUPAC letters minus the four standard bases) plus one extra letter was constructed. The occurrences of the extra letter were then replaced by the standard DNA bases in a cyclic manner (Figure S11 and Table S3**Error! Reference source not found.**).

This 192 base oligo (de Bruijn + primers) was then synthesized, processed and sequenced using similar procedures to the above with the following differences:

- The oligo was synthesized using IDT Ultramer synthesis technology for long synthetic DNA oligos
- Sequencing was performed using the Nano Mi-Seq kit yielding 150bp paired end reads.

We obtained 1,086,991 150bp paired-end reads. We merged the read pairs using PEAR (*14*) to generate 1,017,813 reads, 90% of which with the designed length of 145 bases. We used a similar pipeline to the one described above to calculate inference rates for each position in the sequence and to investigate the properties of the error rates.

### Code availability

All custom code used in this study is available online at:

- https://github.com/leon-anavy/Reed-Solomon Implementation of the custom Reed Solomon encoding and decoding pipeline
- https://github.com/leon-anavy/composite-DNA Implementation of the simulations and analysis results
- https://github.com/leon-anavy/dna-fountain An alteration of the previously published DNA fountain^9^ code

### Data availability

Raw sequencing data that support the findings presented in Table 1 and Figures 3–5 have been deposited in NCBI SRA database with accession codes [TBD].

## Acknowledgments

We thank Tal Katz-Ezov and Tamar Hashimshony from the Technion Genome Center for the advice and assistance with oligo design and sequencing experiments. We also thank Patrick Weiss from TWIST bioscience for technical support and assistance with DNA synthesis. Finally, we thank the Yakhini and Amit research groups for valuable comments and discussions. L.A. is supported by the Adams Fellowships Program of the Israel Academy of Sciences and Humanities. This project received funding from the European Union’s Horizon 2020 Research and Innovation Programme under grant agreement no. 664918—MRG-GRammar

